# ORMDL1 is upregulated and associated with favorable outcome in colorectal cancer

**DOI:** 10.1101/2020.12.10.420497

**Authors:** Qian Wang, Wanjun Liu, Si Chen, Qianxin Luo, Yichen Li, Shaoyong Peng, Huaiming Wang, Xiaoxia Liu, Daici Chen

## Abstract

**Background:** ORMDL1 gene encodes a transmembrane protein for endoplasmic reticulum and is known as crucial negative regulator for sphingolipid biogenesis. However, it has been rarely studied in tumor-related context. Therefore, its prognostic value and functional significance in colorectal cancer (CRC) remain to be explored.

**Methods:** TCGA CRC cohort analysis, qRT-PCR, and immunohistochemistry (IHC) were used to examine the ORMDL1 expression level. The association between ORMDL1 expression and various clinical characteristics were analyzed by Chi-square tests. CRC patients’ overall survival (OS) was analyzed by Kaplan-Meier analysis. *In vitro* and *in vivo* cell-based assays were performed to explore the role of ORMDL1 in cell proliferation, invasion and migration. Transcriptional changes of cells either with ORMDL1 knockdowned or overexpressed were compared and analyzed.

**Results:** ORMDL1 was upregulated in CRC tissues either in TCGA cohort or in our cohort. Interestingly, its expression was significantly lower in patients with metastasis compared to patients without metastasis, and high expression group had longer OS than low expression group. Knockdown of ORMDL1 expression can promote proliferation, colony formation and invasion, while attenuate migration in CRC cell lines. In opposite, forced overexpression of ORMDL1 reduced cell proliferation, colony formation and invasion, while enhanced cell migration. Epithelial-to-mesenchymal transition (EMT) related genes were enriched among differentially expressed genes when ORMDL1 was knockdowned in cells, which was consistent with morphologic change by microscopy observation. Finally, stable knockdown of ORMDL1 can promote cancer cell proliferation *in vivo* to some extent.

**Conclusion:** ORMDL1 is upregulated and may serve as biomarker to predict favourable outcome in colorectal cancer.

## Introduction

Colorectal cancer (CRC) is a malignant tumor with high incience in the gastrointestinal tract, leading to the third most common cancer as well as the second leading cause of cancer mortality overall (1, 2). Usually the occurrence and development of CRC go through four successive stages: initiation, promotion, progression and metastasis, and liver is the most common metastatic organ, followed by the lungs, bones and peritoneum (3). The 5-years survival rate of advanced CRC patients with metastasis to distant organ is very low (below 5%) (4). Although new treatment strategies for CRC besides surgical resection, *e. g.*, radiotherapy, chemotherapy, immunotherapy or targeted therapy have been developed, about 50% of CRC patients who developed metastasis still exhibited poor survival rate (5–7). Some biomarkers had been found to achieve early prognosis as well as for targeted therapy and molecular stratification in CRC (8, 9).Similarly, to find more promising biomarkers for the detection and diagnosis of early metastases of CRC is an valuable and advantageous approach to improve patients’s survival.

Tumor cells possess many unique hallmarks, of which the most important is their ability to proliferate indefinitely, invade and metastasize. Through epithelial–mesenchymal transition (EMT) and remoding of cytoskeleton, certain cancer cells become motile so that they aquire the abilities to migrate and spread to other tissues (10, 11). Among this process, Rho GTPases act as an molecular switch which regulated by guanine nucleotide exchange factors (GEFs) and GTPase-activating proteins (GAPs), to regulate tumor cell morphology, cell adhesion to extracellular matrix and cytoskeleton reorganization (12, 13). Rho GTPases also regulate cell movement by activating a variety of downstream effectors(14, 15). Given the close relationship between Rho GTPases and cancer cells migration and metastasis, targeting Rho GTPases may help prevent tumor from deteriorating(16–18). Moreover, if drugs can block the interaction between Rho GTPase and downstream effectors very specifically in cancer, side effects would become much less than Rho GTPase inhibition in bulk (16). This scenario is also the ideal principle for halting EMT in cancer metastasis.

The endoplasmic reticulum (ER) is essential for the synthesis of proteins and lipids. Studies have showed that endoplasmic reticulum stress is closely related to the occurrence of various diseases, including cancer(19). To induce endoplasmic reticulum stress with cytotoxic drugs can accelerate the apoptosis of cancer cells (20).These indicated that endoplasmic reticulum plays important role in tumor development and progression. ORMDL family (ORMDL1, ORMDL2, ORMDL3) can encode transmembrane protein of endoplasmic reticulum and functions crucially in sphingolipid homeostasis (21, 22), which is regulated by ceramide level sensing (23, 24). It also participates in proper folding of endoplasmic reticulum proteins (25). Related to cancer, it was reported that ORMDL family protein interacts with SPTLC1 and was associated cell renal cell carcinoma development (26). However, the expression of ORMDL1 in CRC and its prognostic significance are unclear, and whether it is involved in the progression of CRC is unknown.

In the current study, we initially investigated The Cancer Genome Atlas (TCGA) cohort and discovered that ORMDL1 was upregulated in CRC tissues. Analysis of clinical characteristics showed that patients with low ORMDL1 expression were more inclined to have metastasis and had shorter OS. By performing various cell-based assays, we investigated the cellular malignant behavior of colorectal caner cells which were influenced by ORMDL1 expression. With the presented evidence, we concluded that ORMDL1 is a valuable and promising biomarker whose hightened expression is associated with favorable outcome in CRC, providing candidate for molecular stratification and potential therapeutic target in CRC comprehensive treatment regimen.

## Materials and methods

### Patients and tissue samples

The frozen paired samples used for mRNA quantification were stocked in RNAlater solution (Invitrogen, Thermo Fisher Scientific, USA), and were acquired from 41 CRC patients with primary CRC tissues and matched adjacent normal tissues, who had undergone surgery between November 2012 and December 2017 at the Sixth Affiliated Hospital of Sun Yat-sen University (SYSU).

We also constructed tissue microarrays of primary CRC from 217 patients from the Sixth Affiliated Hospital of SYSU. None of these patients received chemotherapy, radiotherapy and other related treatment prior to surgical resection of the tumor. Among these patients, 122 patients had disease relapse or metastasis within 3 years after surgery. Our study was approved by the Institutional Ethics Committee of Sixth Affiliated Hospital, Sun Yat-sen University.

### Cell culture and transfection

All cell lines (HCT15, HCT8, SW480, SW620, LOVO, DLD1, SW48, HCT116, 293T) used in this study were obtained from ATCC. Cell lines were typically cultured at 37℃ in an incubator with 5% CO2. SiRNAs used were as follows: siORMDL1#1: 5’-GGAGTTGGCTTGCTTCATA-3’, siORMDL1#2: 5’-CTGGCAAGTTTCTATACGA-3’. Negative scramble fragment (RiboBio Co., China) were used as control.

### Quantitative real time PCR (qRT-PCR)

The mRNA was extracted from CRC cell lines and tissues with TRIzol (Invitrogen, CA, USA). The sequence of the PCR primers were as follows: ORMDL1, forward: 5’-TTC AGT GTT CCT GTT GCT TG-3’and reverse: 5’-AGC CTT GCT TTA CCC TGG TC-3’; β-actin, forward: 5′-TTG TTA CAG GAA GTC CCT TGC C-3′,and reverse 5′-TTG TTA CAG GAA GTC CCT TGC C-3′. β-actin was used for normalization. The siRNAs were transfected into CRC cells via Lipofectamine RNAiMAX (Invitrogen) and overexpression plasmids of ORMDL1 (Full length ORMDL1 cDNA inserted onto pEZ-M98 plasmid, FulenGen, Guangzhou, China) were transfected with Lipofection 3000 (Invitrogen).

### Western blotting

Cells were lysed and boiled for 10 minutes. Proteins were separated by SDS-PAGE, transferred to PVDF membrane, and detected with relevant antibodies for RhoA (1:1000 dilution, Cat#2117, Cell signaling technology), RhoB (1:1000 dilution, Cat#63876, Cell signaling technology), RhoC (1:1000 dilution, Cat#3430, Cell signaling technology), Slug (1:1000 dilution, Cat#9585, Cell signaling technology), Snail (1:1000 dilution, Cat#3879, Cell signaling technology), and ZEB1 (1:1000 dilution, Cat#3396, Cell signaling technology). Blotting for α-tubulin (1:20000, Cat#66031-1-Ig, Proteintech) was used as a loading control.

### Hematoxylin and eosin (H&E) and immunohistochemistry assay

The tissue microarray and related clinicopathologic information were collected from Sixth Affiliated Hospital of SYSU. IHC for ORMDL1 was carried on the CRC tissue microarray slides. The slides were first incubated in 60℃ for 4h following deparaffinized in xylene, then rehydrated in alcohol. After heating in citrate buffer for 21 minutes, we used the IHC kit (cat# SP9000; ZSGB-Bio, Beijing, China) to block the endogenous peroxidase activity. Slides were bloked with 5% goat serum for 1hour (h), and incubated in the anti-ORMDL1 antibody (1:100, PHA065643, Atlas Antibodies) overnight at 4℃. Next day, after 3 times washed by PBST, slides were incubated with secondary antibody for 1h, then stained by DAB kit (cat# ZLT-9017; ZSGB-Bio). After experiments, slides were observed by microscope. The IHC scores were independently assessed by two pathologists. And IHC score of 6.3 was selected as the cutoff value to divide the high and low expression of ORMDL1.

### Cell proliferation assay

HCT116 and DLD1 were selected to perform this experiment. We separately seeded 5000 DLD1 cells and HCT116 cells after transfection with ORMDL1 overexpress plasmid or siORMDL1 into 96-well plates. The real-time cell analyzer (RTCA, xCELLigence system, ACEA Biosciences, Inc.) and IncuCyte Essens Bioscience incubator (Essens Bioscience, Birmingham, UK) were used to monitor cell growth. The live cells were recorded automatically every 30 minutes.

### Colony formation assay

500 cells were seeded into 6-well plate, and incubated for 8 days at 37℃ incubator. Then colonies were fixed with 4% paraformaldehyde and stained with 0.1% crystal violet.

### Wound-healing assay

After transfection, cells were seeded into 24-well plates and incubated until 100% confluence. A scratch was made in the cell layer using 10ul pipette tip, then HCT116 were cultured in FBS-free McCoy’s 5A and DLD1 in FBS-free RPMI-1640. Live cell photos were collected by the IncuCyte Essens Bioscience incubator every 30 minutes.

### Invasion assays

8-μm transwell filters (Coring, NY, USA) were used to perform the invasion assay. The upper chambers were covered with Matrigel (Corning, NY, USA), and placed in the 37℃ incubator for 1h. Then 8×10^4^ DLD1 cells were seeded into the upper chamber with RPMI-1640 and we added 700ul RPMI-1640 with 10% FBS to the 24-well plates. After 24h, we wiped the upper chamber with cotton swabs, and stained cells in the other side of the membrane with crystal violet for 5min. The other cell line HCT116 was conducted with the same method. But it was cultured with McCoy’s 5A medium and was planted for 10×10^4^ cells.

### *In vivo* subcutaneous tumor growth assay and H&E and IHC assay

The shORMDL1 plasmid (ORMDL1 specific shRNA sequence same as siORMDL1#1 inserted onto pLKD-CMVEGFP-2APuro-U6-shRNA plasmid, Obio Tech, Shanghai, China) or empty control plasmid were stably transfected into DLD1 cells used lentivirus transfection method, and subjected to stable selection by puromycin. The BALB/c nude mice were injected with 4×10^6^ stable shORMDL1 DLD1 cells in each mouse subcutaneously, and each group had 7 mice. Tumor sizes were measured every 3 days then calculated as volume = 1/2 (length x width^2^). After 15 days, mice were scarified and tumors were measured then collected. All specimens were fixed by formalin and embedded by paraffin for slide preparation. Slides were stained by H&E and anti-Ki67 accordingly.

### Immunofluorescence (IF) staining

DLD1 cells layered on confocal-compatible slides were transfected with control or siORMDL1 reagent, then fixed 24 hours afterwards with 4% paraformaldehyde. After being incubated with anti-ORMDL1 antibody (1:100, PHA065643, Atlas Antibodies) and following secondary antibody according to product manual, all slides were observed and photoed (Leica SP8, Mannheim, Baden-Wuerttemberg, Germany).

### Statistical analysis

All statistical analysis was performed using GraphPad Prism 8.0 (Chicago, IL, USA) or SPSS 25.0 (California, USA). Association between ORMDL1 expression and clinical variables were analyzed by Chi-square tests. Overall survival was analyzed using Kaplan-Meier analysis and the *p* value was calculated by log-rank test. COX regression analyses model was used to evaluate univariate and multivariate survival analyses. Differences between groups were analyzed by a two-tailed Student’s t-test or a Mann-Whitney U-test.

### RNA sequencing

HCT116 cells with transilent transfection of either ORMDL1 overexpression plasmid or knockdown (siORMDl1 and siControl) reagent, were collected 48 hours after transfection. Cell medium was collected as well for contamination detection for mycoplasma. Whole transcriptome deep sequencing (RNA seq) was conducted by GeneSeed Company (Guangzhou, China) with HiSeq X10 PE150 platform. Differential genes were generally defined as the difference between the two groups more than 1.5 times and *p<0.05*.

## Results

### ORMDL1 expression was increased in CRC and associated with favorable outcome

Analysis of TCGA data (50 CRC cases with self peri-tumor tissue matched) showed that the mRNA level of ORMDL1 was obviously higher in tumor than in peri-tumor tissues (Fig 1a). We also examined the ORMDL1 mRNA level in primary tumor, peri-tumor (~2cm from tumor), normal tissues (~5cm from tumor) and metastatic tumor among 41 CRC patients independetely. We found that ORMDL1 expression was increased in CRC primary tumors against peri-tumor tissues and normal tissues (Fig 1B). Moreover, it expressed significantly higher in the metastatic tissues (Fig 1b).

**Figure 1:**
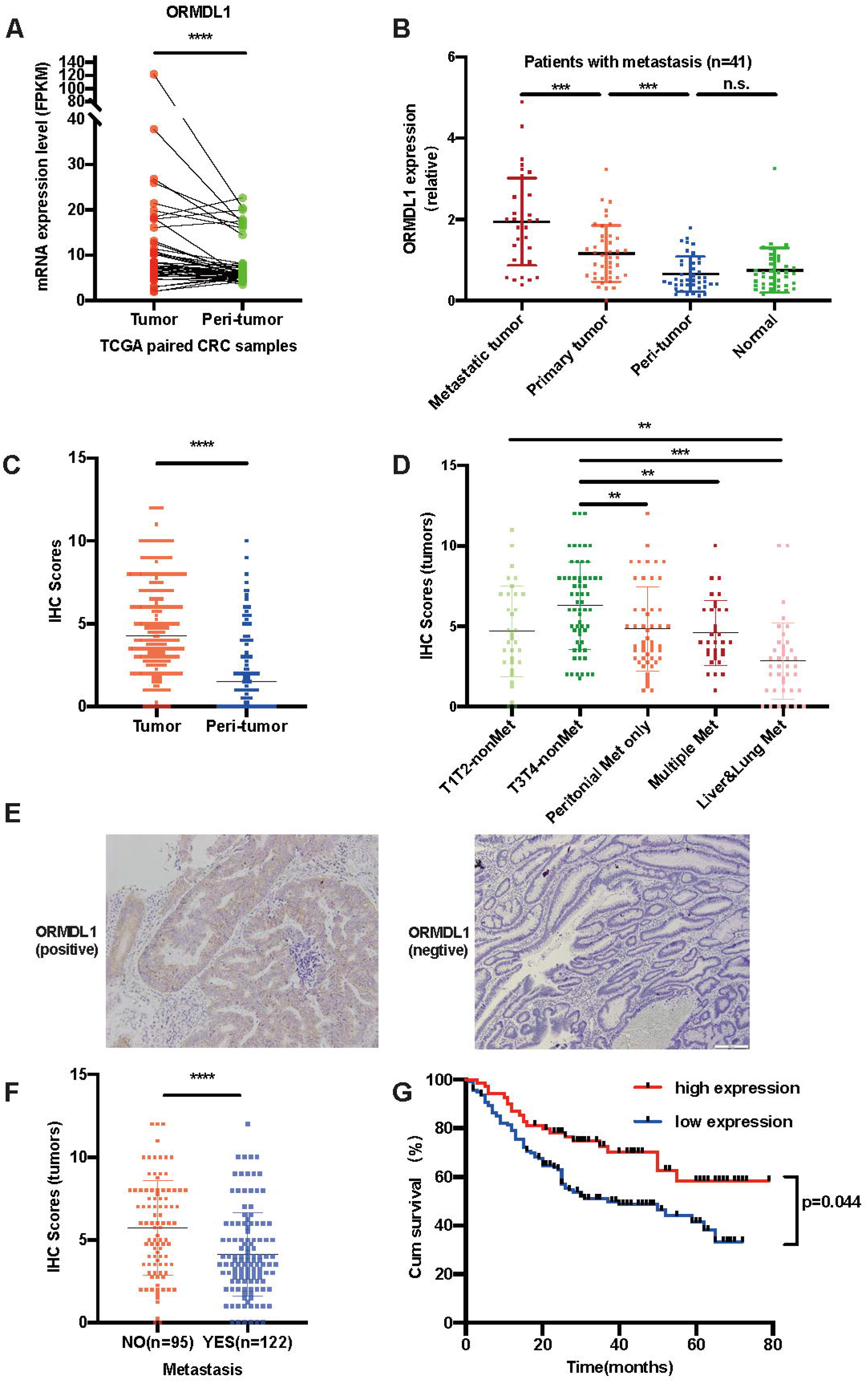
ORMDL1 expression was increased in CRC and associated with favorable outcome. (a) The mRNA expression level of ORMDL1 was significantly higher in CRC tissues compared with peri-tumor tissues in TCGA cohorts. n=50, ****p<0.0001, two-tailed Student’s t-test. (b) The mRNA expression level of ORMDL1 in CRC primary tumors, matched peri-tumor, metastatic tumor and normal tissues of patients. ***p<0.001, ns not significant (p>0.05), two-tailed Student’s t-test. (c) IHC scores of ORMDL1 expression level in tissue microarray of tumor and peri-tumor tissues from patients. ****p<0.0001, two-tailed Student’s t-test. (d) IHC scores of ORMDL1 expression level in tissue microarray of T1/T2-nonmetastasis group, T3/T4-nonmetastasis group, peritoneal metastasis group, liver and lung metastasis grpup and multiple metastasis group. **p<0.01, ***p<0.001, two-tailed Student’s t-test. (e) IHC staining images of ORMDL1 in tissue microarray of paired samples of CRC tissues (left image) and adjacent tissues (right image). (f) IHC scores of ORMDL1 expression level in tissue microarray of patients with or without metastasis. ****p<0.0001, two-tailed Student’s t-test. (g) Kaplan-Meier survival curves of overall survival based on IHC scores of ORMDL1 expression in tissue microarray. P=0.0044, log-rank test.

To further investigate the association between ORMDL1 expression and patients’ clinicopathologic characteristics, we performed IHC analysis based on the paraffin-embedded tissue microarray of 217 CRC patients in our center. The ORMDL1 positive staining mainly concentrated in cell cytoplasm of CRC tissues; on the contrary, the adjacent normal glands presented negative or low level (Fig 1c). IHC scoring revealed that similar to the ORMDL1 mRNA level, ORMDL1 protein level was upregulated in tumor compared to the adjacent tissues (Fig 1d). IHC scores were lower in the metastatic patients, but different metastatic groups expressed different level of ORMDL1 (Fig 1e). Among the cohort, patients with metastasis were significantly lower than those without metastasis (Fig 1f). According to the ROC curve analysis, we selected the IHC score of 6.3 as the cutoff value which can best separated the cohort into two groups: high (n=69) and low (n=141) ORMDL1 expression. Among all the clinical characteristics analyzed, we found that the depth of invasion, distant metastasis, TNM stage and nerve invasion were related to the ORMDL1 expression (Table 1). The multivariate analysis showed that ORMDL1 expression was not an independent predictor for CRC patients’ OS (Table 2). Kaplan-Meier analysis showed that the high expression group had better overall survival (p=0.004) (Fig 1g).

**Table 1.**
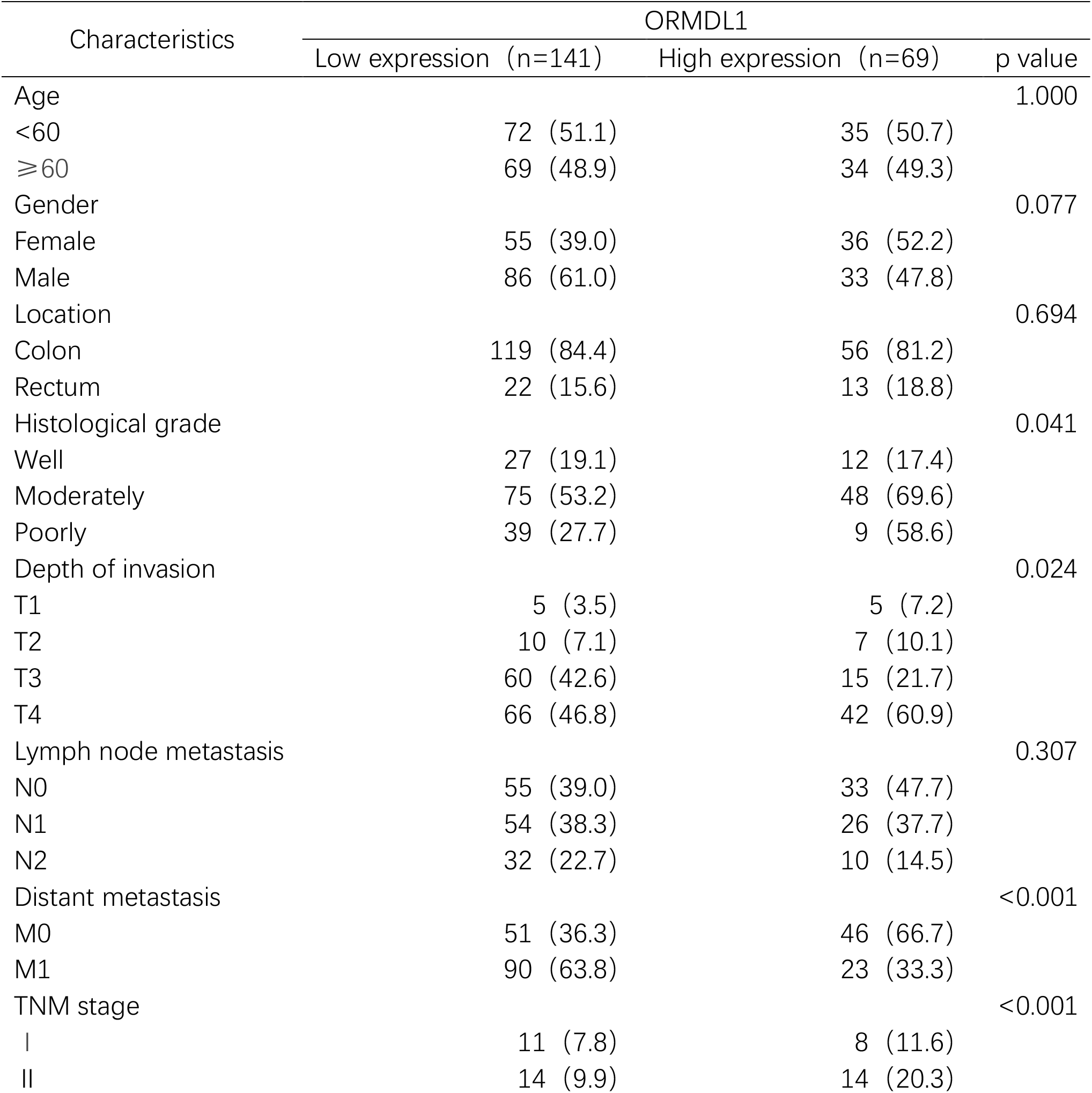
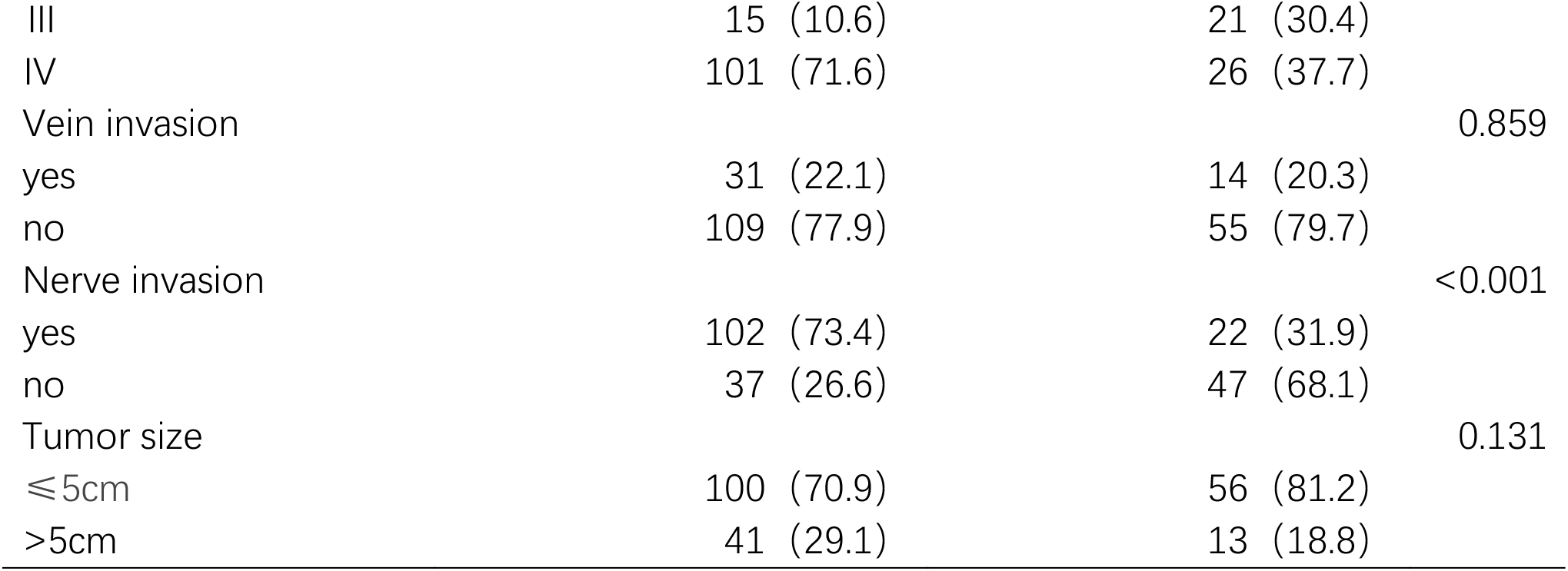
Correlation between expression of ORMDL1 and clinicopathological features of CRC patients.

**Tabel 2.**
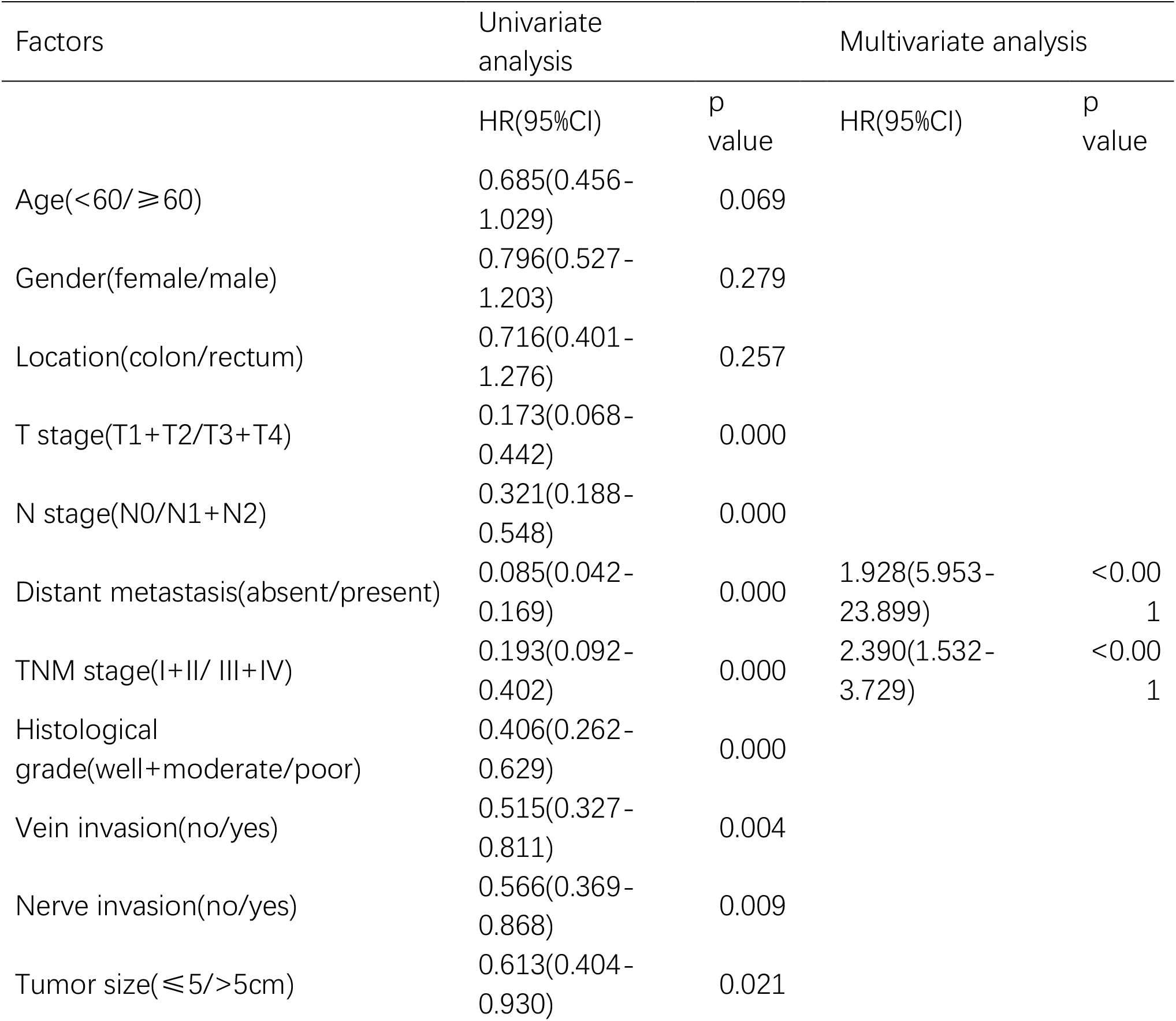
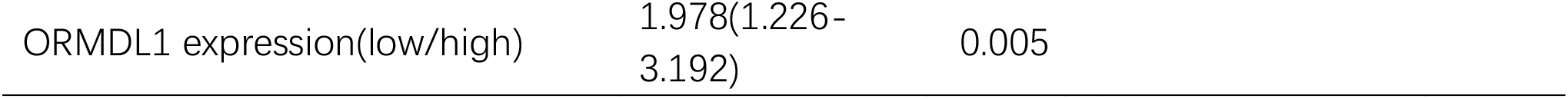
Univariate and multivariate analysis of different prognostic parameters of the CRC patients in the cohort.

| Factors | Univariate analysis |  | Multivariate analysis |  |
| --- | --- | --- | --- | --- |
|  | HR(95%CI) | p value | HR(95%CI) | p value |
| Age(<60/≥60) | 0.685(0.456-1.029) | 0.069 |  |  |
| Gender(female/male) | 0.796(0.527-1.203) | 0.279 |  |  |
| Location(colon/rectum) | 0.716(0.401-1.276) | 0.257 |  |  |
| T stage(T1+T2/T3+T4) | 0.173(0.068-0.442) | 0.000 |  |  |
| N stage(N0/N1+N2) | 0.321(0.188-0.548) | 0.000 |  |  |
| Distant metastasis(absent/present) | 0.085(0.042-0.169) | 0.000 | 1.928(5.953-23.899) | <0.001 |
| TNM stage(I+II/ III+IV) | 0.193(0.092-0.402) | 0.000 | 2.390(1.532-3.729) | <0.001 |
| Histological grade(well+moderate/poor) | 0.406(0.262-0.629) | 0.000 |  |  |
| Vein invasion(no/yes) | 0.515(0.327-0.811) | 0.004 |  |  |
| Nerve invasion(no/yes) | 0.566(0.369-0.868) | 0.009 |  |  |
| Tumor size(≤5/>5cm) | 0.613(0.404-0.930) | 0.021 |  |  |

|  |  |  |
| --- | --- | --- |
| ORMDL1 expression(low/high) | 1.978(1.226-3.192) | 0.005 |

### Knockdown of ORMDL1 expression promoted proliferation and invasion but inhibited migration of CRC cells

To explore the potential role of ORMDL1 gene in CRC cellular behavior, we first estimated the ORMDL1 expression level in 8 CRC cell lines and a human kidney epithelial cell line 293T, and discovered that ORMDL1 expressed relatively low in all CRC cell lines compared to 293T cells (Fig 2a). HCT116 and DLD1 were chose for all following cell-based experiments. ORMDL1 was successfully silenced in HCT116 and DLD1 by siRNA transfection (Fig 2b). Real time monitoring of cell growth by RTCA indicated that knockdown of ORMDL1 expression promoted HCT116 proliferation (Fig 2c). Colony formation assay also confirmed that ORMDL1 knockdown increased colony numbers, either in HCT116 cells (Fig 2d) or DLD1 cells (Fig 2e). Interestingly, while knockdown of ORMDL1 enhanced the invasion behavior of CRC cells (Fig 2f, 2g), it inhibited the migration capability of CRC cells (Fig 2h, 2i).

**Figure 2:**
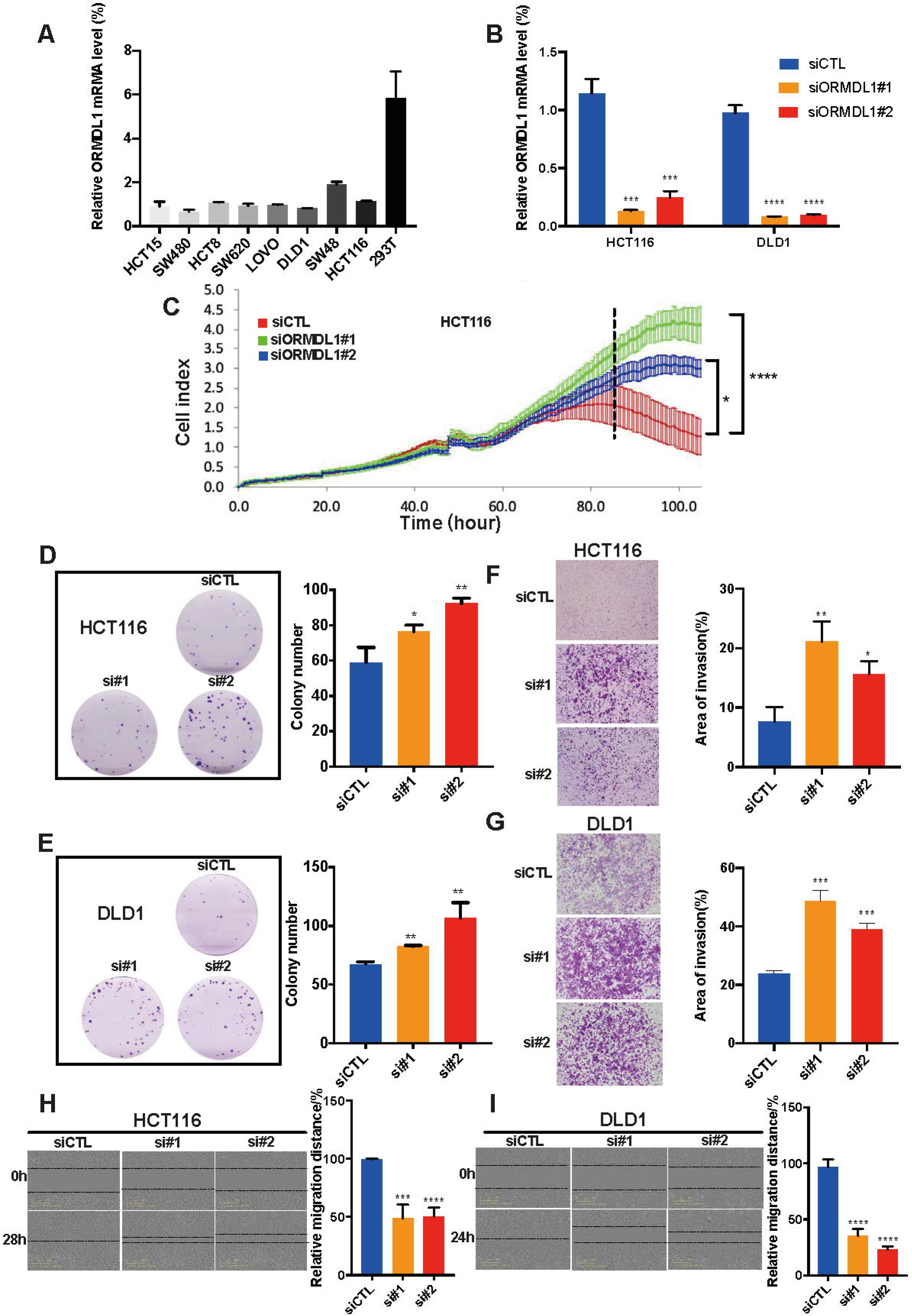
Knockdown of ORMDL1 promoted proliferation and invasion but inhibited migration of CRC cells. (a) Relative mRNA expression level of ORMDL1 in human cell lines was analyzed by qRT-PCR. (b) Knockdown efficiency of ORMDL1 in HCT116 and DLD1 was analyzed by qRT-PCR. ***p<0.001, ****p<0.0001, two-tailed Student’s t-test. (c) Proliferation assay by RTCA showed that downregulation of ORMDL1 can promote the proliferation ability in HCT116. ****p<0.0001, *p<0.05, two-tailed Student’s t-test. (d-e) Images and statistical analysis of colony formation assays in two cell lines. *p<0.05, **p<0.01, two-tailed Student’s t-test. (f-g) Images and statistical analysis of wound invasion assays in two cell lines. *p<0.05, **p<0.01, ***p<0.001. (h-i) Images and statistical analysis of wound healing assays in two cell lines. ***p<0.001, ****p<0.0001.

### Overexpression of ORMDL1 attenuated proliferation and invasion but promoted migration of CRC cells

In order to further determine the effect of ORMDL1 expression on CRC cells, we transfected HCT116 and DLD1 with an ORMDL1 overexpression vector; the overexpression level of ORMDL1 was confirmed (Fig 3a). Proliferation of CRC cells seemed to be suppressed by ORMDL1 overexpression (Fig 3b), consistent with results from colony formation assay (Fig 3c). Interestingly, while overexpression of ORMDL1 attenuated the invasion behavior of CRC cells (Fig 3d), it promoted the migration capability of CRC cells (Fig 3e). These results suggest that ORMDL1 expression plays non uniform roles in the proliferation, migration and invasion behavior of CRC cells.

**Figure 3:**
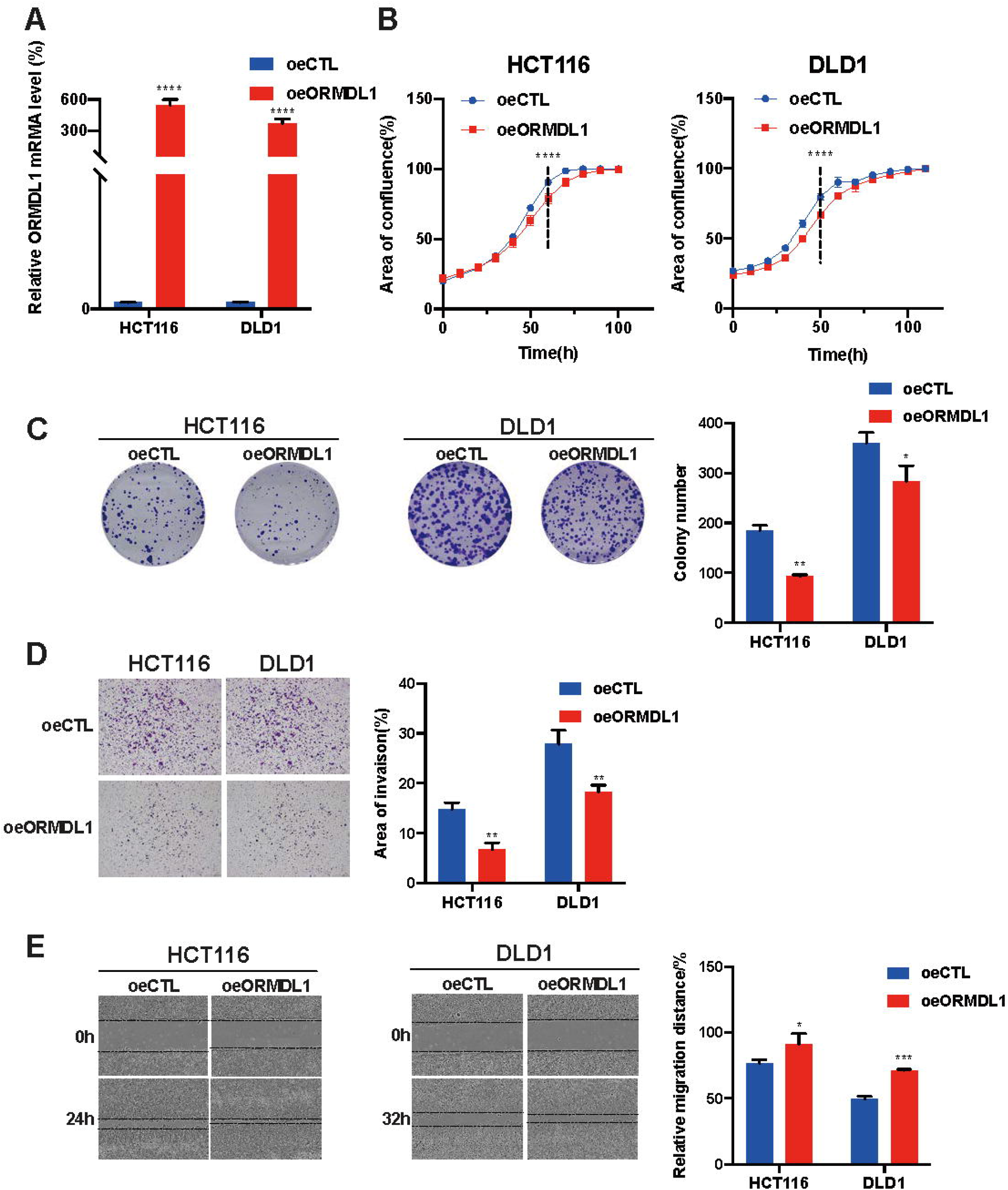
Overexpression of ORMDL1 attenuated proliferation and invasion but promoted migration of CRC cells. (a) Overexpression of ORMDL1 in HCT116 and DLD1 was analyzed by qRT-PCR. ****p<0.0001, two-tailed Student’s t-test. (b) Proliferation assays recorded by IncuCyte showed that overexpression of ORMDL1 promoted proliferation in CRC cells. (c) Images and statistical analysis of colony formation assays showed that overexpression of ORMDL1 inhibited the colony formation ability in CRC cells. *p<0.05, **p<0.01, two-tailed Student’s t-test. (d) Images and statistical analysis of invasion assays indicated that overexpression of ORMDL1 inhibited CRC cells invasion. **p<0.01, two-tailed Student’s t-test. (e) Images and statistical analysis of wound-healing assays showed that overexpression of ORMDL1 can promote the migration ability in CRC cells. *p<0.05, ***p<0.001, two-tailed Student’s t-test.

### ORMDL1 modulates cytoskeleton remodeling potentially through affecting expression of EMT-related genes

ORMDL1 is known to be membrane-bound on the endoplasmic reticulum. We hypothezed that cell movement related genes might be affected by ORMDL1 expression. To explore potential mechanism through which ORMDL1 excecutes its function during the progression of CRC, we performed transcriptome RNAseq analysis in cells with ORMDL1 knockdowned or overexpressed. By filtering for significant expressed genes (|fold change| >= 1.5, *p < 0.05*, *FDR < 0.05*), we identified 76 differential expressed genes (DEGs) in the overexpression group (oeORMDL1 *vs* oeControl, Supplemental Table 1) and 2329 DEGs in the knockdown group (siORMDL1 *vs* siControl, Supplemental Table 2). As the DEGs list in overexpression group was too small to be subjected to enrichment analysis (usually 10% of the whole transcriptome), we only focused on DEGs in the knockdown group in following analysis, which included 1222 up-regulated genes and 1107 down-regulated genes (Volcano plot, Fig 4a). Within the DEGs of the knockdown group comparison, EMT, TRAIL signaling and integrin signaling were showed as most significant enriched pathways for the upregulated genes (Fig 4b). We also confirmed some of the most DEGs by qRT-PCR (Fig 4c) and Western bloting (Fig 4d). For example, some EMT-related transcriptional factors, Snail, Slug and ZEB1 were affected by ORMDL1 to different extents. Interestingly, the best known Rho GTPases, RhoA, RhoB and RhoC, were also influenced by ORMDL1 expression. Phenotypically, when transfected with siORMDL1 reagent, manifested flat, long spindle shaped and intended to be gathered cells would transform into mesenchymal cell morphology: uniformly round shape with clear boundaries (Fig 4e). This dramatic morphologic change indicated that EMT was induced in the cells.

**Figure 4:**
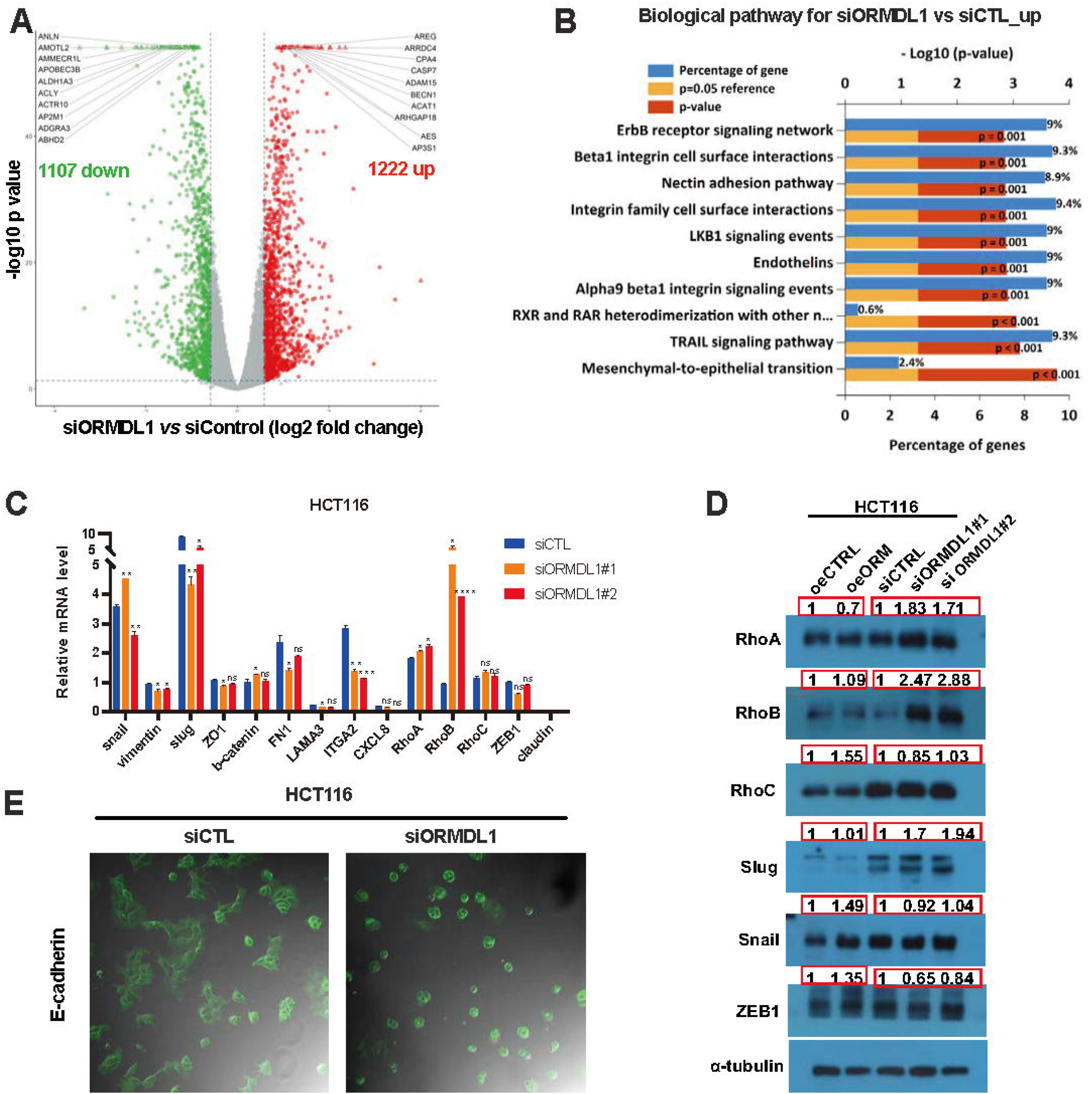
ORMDL1 modulates cytoskeleton remodeling potentially through affecting expression of EMT-related genes. (a) Volcano plot for genes identified by RNA sequencing, with X axis for the log2 fold change (siORMDL1 *vs* siControl) and Y axis for -log10 p value. (b) List of upregulated genes in (a) was subjected to pathway enrichment analysis and the top 10 enriched pathways is showed. (c) qRT-PCR to confirm DEGs whose expression level was changed when ORMDL1 was knockdowned in HCT116 cells. (d) Western blotting and semi-quantitative analysis for Rho family genes and EMT-related genes when ORMDL1 was knockdowned or overexpressed in HCT116 cells. (e) E-cadherin IF overlayed with bright field image for cells transfected with siCTL or siORMDL1.

### Knockdown of ORMDL1 expression can promote tumor growth *in vivo*

Based on the clinicopathologic analysis and *in vitro* assays above, we continued to further investigate whether knockdown of ORMDL1 affects tumor growth *in vivo* by performing tumor-forming experiments in nude mice. Tumor volumes were recorded every 3 days and results showed that knockdown of ORMDL1 (shORMDL1) promoted the proliferation of cancer cells *in vivo* to some extent (Fig 5a). The shORMDL1 group showed larger tumor formation, but without statistical significance in term of tumor weights (Fig 5b). We used qRT-PCR assay to confirm that ORMDL1 was successfully knockdowned in the tumor cells (Fig 5c). HE staining showed that the dissected tumors were indeed tumorous and not inflammatory tissue (Fig 5d, upper panel). IHC staining for Ki67 showed that more intensive Ki67 signals in the shORMDL1 group (Fig 5d, lower panel and quantification, Fig 5e), suggesting cancer cells were proliferating vigorously to form a tumor.

**Figure 5:**
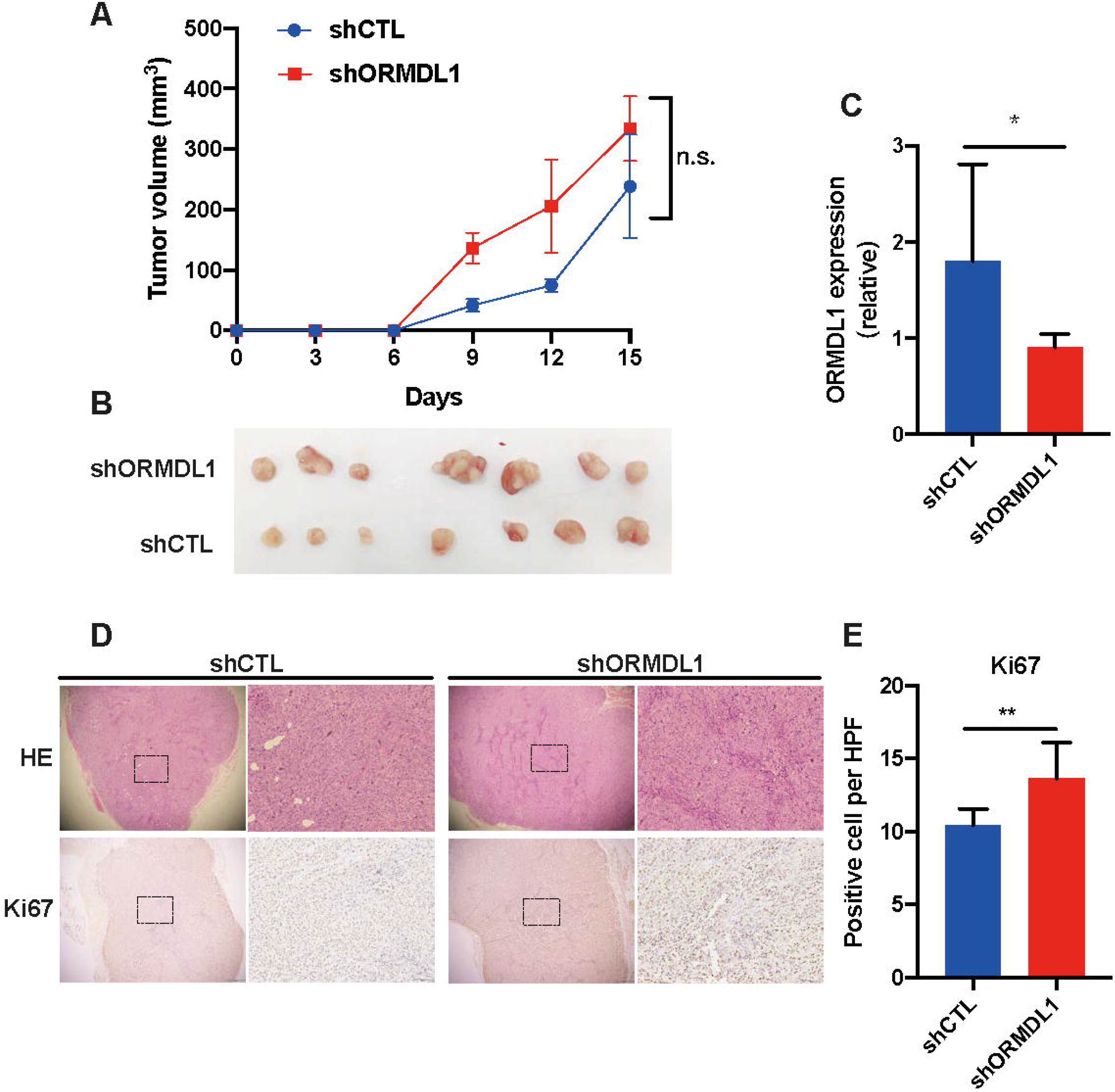
Knockdown of ORMDL1 expression can promote tumor growth *in vivo*. (a) ORMDL1 stably knockdowned cells were subcutaneously implanted for tumor model in nude mouse as described in Methods. Tumor volumes were recorded and calculated (n=7). (b) After 15 days, mice were scarified and tumors were collected and weighted. (c) Dissected tumor from mice were subjected to qRT-PCR analysis for ORMDL1 level. (d) Dissected tumors were subjected to HE staining and Ki67 IHC staining (e) Ki67 positive cells per 20x high power field (HPF) were calculated.

## Discussion

Human ORMDL gene family has three members ORMDL1, ORMDL2 and ORMDL3, all encoding transmembrane proteins anchored in the endoplasmic reticulum (25). This gene family seems to be very conserved across eukaroytic cells, from singled cell budding yeast to mammalian cells(25). Genome-wide association studies (GWAS) showed that there were significant association between mutation in ORMDL genes (ORMDL3 in particular) and asthma(27). Later on ORMDL3 was identified as an inducible lung epithelial gene regulating metalloproteases, chemokines, OAS, and ATF6, establishing the milestone molecular basics for its functions in allergic response(28). Another milestone finding was that proteins of the ORMDL family were shown to be negative homeostasis regulators of serine palmitoyl transferase (SPT), which is the rate-limiting enzyme for the biosynthesis of new sphingolipids (29, 30). Within these three genes, ORMDL3 and ORMDL1 seem to be prevalent players either in allergic response or sphingolipids regulation. Although genomic locus of ORMDL3 has been revealed to be associated with childhood acute lymphoblastic leukemia by GWAS(31), little was known for their roles in the context of other disease, *i.e.*, human cancer. ORMDL3 was reported to be involved in triple-negative breast cancer cell proliferation regulated by upstream factors long noncoding RNA SOX21-AS1/miR-520a-5p(32). Yet, the direct link between ORMDL and cancer progression is to be explored.

In our study, by examing the expression level of ORMDL1 mRNA in TCGA cohort as well as in our cohort, we found that ORMDL1 was obviously upregulated in CRC tumor tissues compared with adjacent normal tissues. In tissues with metastasis groups, ORMDL1 was downregulated compared with tissues without metastasis according to IHC staining scores. Among all the clinical characteristics analyzed, we found that the depth of invasion, distant metastasis, TNM stage and nerve invasion were associated with ORMDL1 low expression. Interestingly, patients with higher expression of ORMDL1 were less likely to develop metastasis, therefore for better OS. These interesting phenomenon all indicated that high expression of ORMDL1 is associated with better clinical outcome so that it could potentially serve as a marker for screening patients with better prognosis. It is worthy investigating whether to combine ORMDL1 expression and other biomarkers (PD-L1, MSI *etc*.) would have better stratification for CRC patients.

This study is only a starting point to bring attention that ORMDL1 function in the context of cancer. Detailed molecular mechanism dissection will shed light on the direct link between ORMDL1 and cancer development, for example, how ORMDL1 regulates EMT and Rho GTPase. In a nutshell, we found that ORMDL1 could promote the migration and suppress the invasion as well as proliferation of CRC cells, suggesting ORMDL1 modulating cancer cell aggressiveness in multiple ways. EMT maybe a direct process that could be influenced by ORMDL1 expression, although non-exclusively for small Rho GTPases also such as RhoA, RhoB and RhoC. These RhoGTPase are curcial upstream effectors for coordinating the remodeling of cytoskeleton in space and time. ORMDL1 is ER membrane-bound protein and the direct molecular link to downstream effectors is to be uncovered.

Despite its critical role in sphinfolipid metablism, it might also affect the survival of cancer cells by coregulating with serine palmitoyltransferase long chain base subunit 1 (SPTC1) in the tumor microenvironment and helping SPTC1 be a tumor suppressor protein in clear cell renal cell carcinoma (ccRCC) (26). Based on the relationship between SPTC1 and ORMDL family, we hypothesized that whether ORMDL1 could also work on CRC progression. Previous studies have showed that autophagy of ORMDL1 takes an important part in regulating free cholesterol (FC) metabolism and stimulating sphingomyelin biosynthesis(21, 33). Human abdominal cavity has a variety of cell components, active substances, etc. together constitute a complex microenvironment, and it also rich in visceral adipose tissue. The triacylglycerol phosphate produced in fat cells can be decomposed into glycerol and free fatty acids by lipase in the body. We guessed this FC-rich microenvironment provides suitable soil for peritoneal transfer of CRC. When FC loaded in CRC cells, ORMDL1 binds to autophagosomes then causes autophagy, resulting in a decrease in ORMDL1 in tumor cells. Thus, we guessed that ORMDL1 may be also an important hub for regulating FC the by influence the ER stress and apoptosis.

It is also worth to mention that very few DEGs were identified when performing transcriptome change for oeORMDL1 vs oeControl. The overexpression level seemed to be reasonable based on qRT-PCR. Unfortunately, we had been not able to detect the protein level by Western blotting, as no commercial monoclonal antibody is available. We can only infer that ORMDL1 was overexpressed by using an ORMDL3 antibody that could recognize ORMDL1-3 potentially(24). As ORMDL1 is a membrane-bound protein, co-factor or receptor inside the cell may be needed for downstream signaling transduction. Also it is tempting to speculate that functionalities of ORMDL1 protein could be regulated by post-translation modification, like phosphorylation or ubiquitination.

## Conclusion

ORMDL1 was upregulated in CRC, and patients with higher ORMDL1 expression had longer OS, suggesting that ORMDL1 expression may serve as biomarker for predicting favorable outcome. ORMDL1 may be involved in EMT regulation and cytoskeleton remodeling in CRC, the molecular mechanism yet to be revealed.

## List of abbreviations

CRC: colorectal cancer
IHC: immunohistochemistry
OS: overall survival
EMT: epithelial-to-mesenchymal transition
GEF: guanine nucleotide exchange factors
GAP: GTPase-activating proteins
TCGA: The Cancer Genome Atlas
HE: hematoxylin and eosin
RTCA: real-time cell analyzer
IF: immunofluorescence
DEG: differential expressed gene
qRT-PCR: quantitative real time PCR
ER: endoplasmic reticulum
GWAS: genome-wide association studies
MSI: microsatellite instability
SPTC1: palmitoyltransferase long chain base subunit 1
ccRCC: clear cell renal cell carcinoma
FC: free cholesterol

## Ethics approval and consent to participate

Our study involving human specimens was approved by the Institutional Ethics Committee of Sixth Affiliated Hospital, Sun Yat-sen University. The related ethical approval code is: 2020ZSLYEC-065. The animal study was approved by the Laboratory Animal Center of The Sixth Affiliated Hospital of Sun Yat-sen University, and approval code is: SYXK(YUE)2018-0190.

## Consent for publication

Not applicable.

## Availability of data and materials

All data generated or analysed during this study are included in this article and its supplementary information files.

## Authors’ contributions

CDC conceived and supervised this study. QW, and WJL performed and analyzed most of the cell-based assays. SC and XXL helped cell culture and animal experiment. YCL performed RNA extraction. QXL, SYP and HMW contributed for patient selection and collection of clinical information. QW frafted the manuscript and CDC edited it. All authors read and approved the final manuscript.

## Acknowledgements

Dr. Lei Wang was one of the pioneers for the Sixth Affiliated Hospital of Sun Yat-sen University. We were deeply sorrowful for the loss of him. We will move on with what he had taught and inspired us.

